## Supplemental materials 1 (Figures and Methods) for "Targeted genomic sequencing with probe capture for discovery and surveillance of coronaviruses in bats"

### SUPPLEMENTAL FIGURES

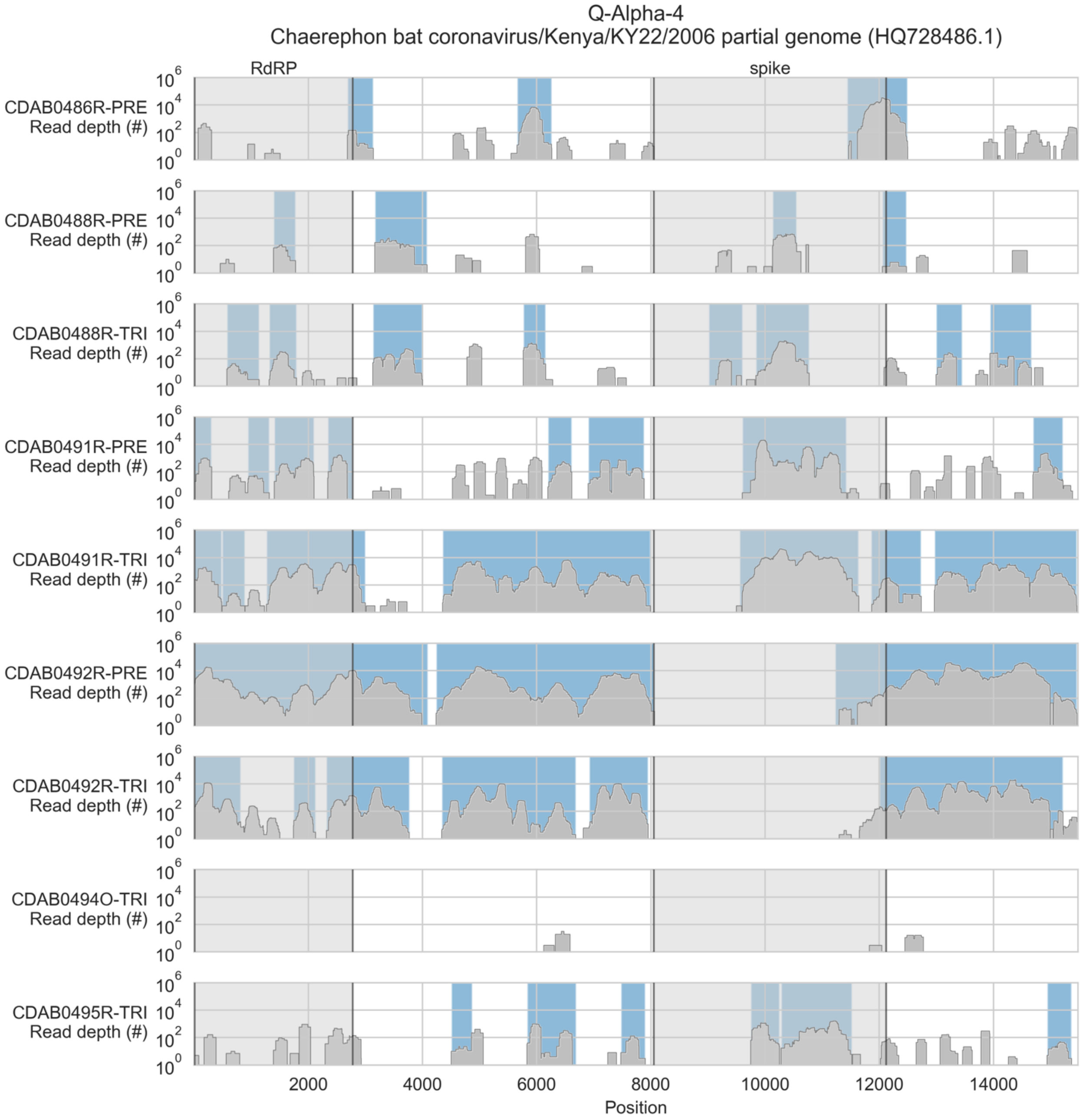


**Figure S1: Coverage of reference sequence by probe captured libraries for specimens from phylogenetic group Q-Alpha-4.** Coverage of reference sequence was determined by mapping reads and aligning contigs from probe captured libraries. Dark grey profiles show depth of read coverage along reference sequence. Blue shading indicates spans where contigs aligned. The locations of spike and RNA-dependent RNA polymerase (RdRP) genes are indicated and shaded light grey.

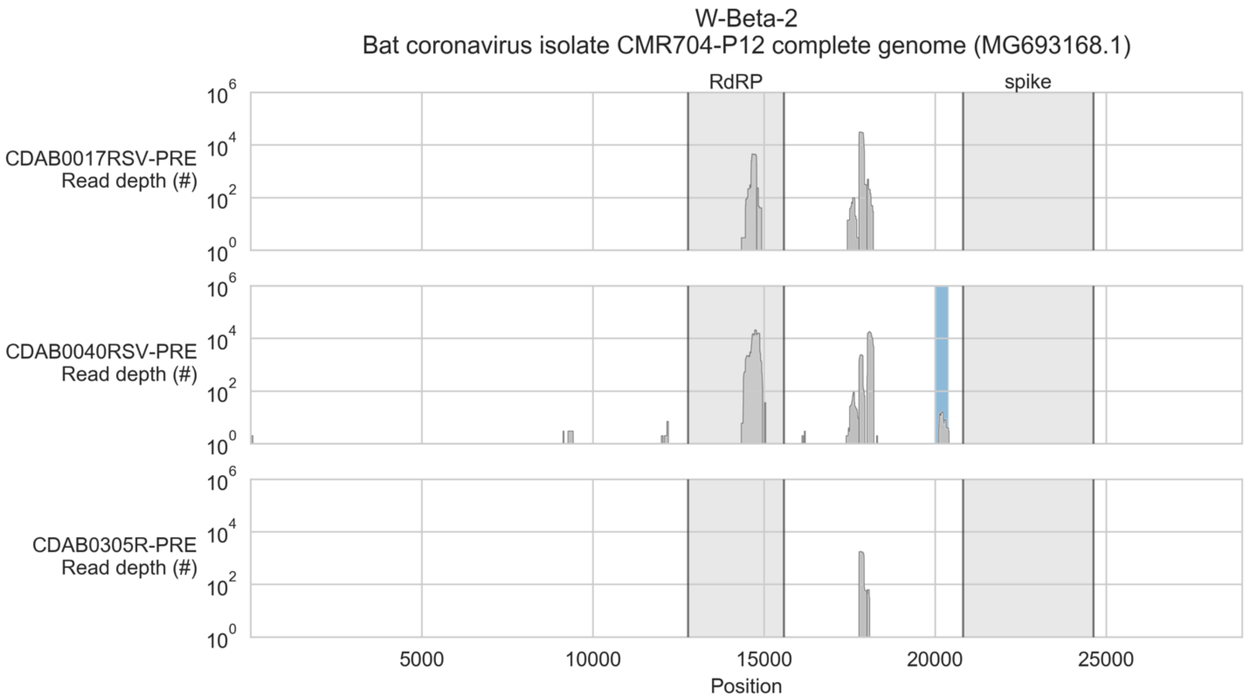


**Figure S2: Coverage of reference sequence by probe captured libraries for specimens from phylogenetic group W-Beta-2.** Coverage of reference sequence was determined by mapping reads and aligning contigs from probe captured libraries. Dark grey profiles show depth of read coverage along reference sequence. Blue shading indicates spans where contigs aligned. The locations of spike and RNA-dependent RNA polymerase (RdRP) genes are indicated and shaded light grey.

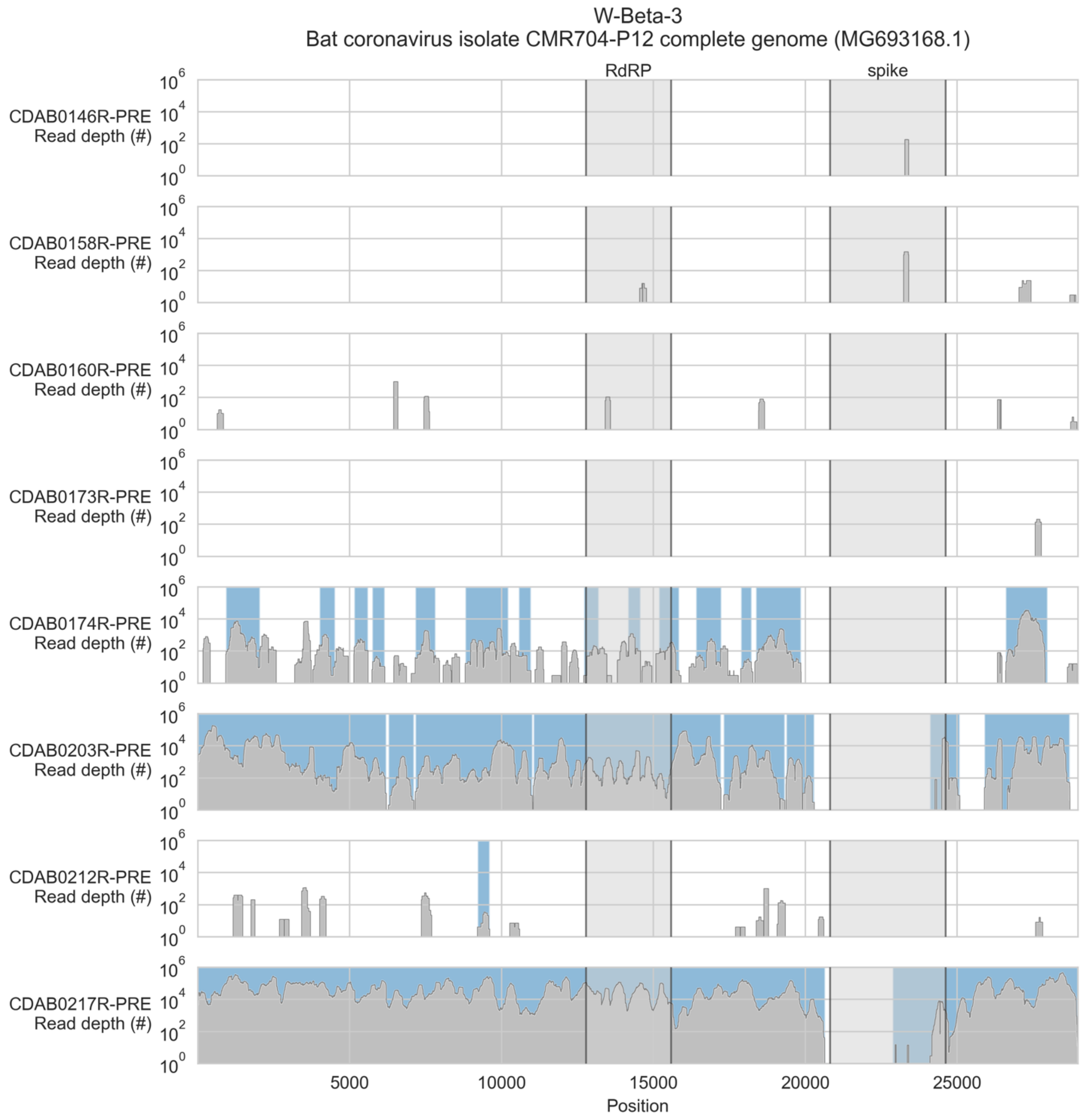


**Figure S3: Coverage of reference sequence by probe captured libraries for specimens from phylogenetic group W-Beta-3.** Coverage of reference sequence was determined by mapping reads and aligning contigs from probe captured libraries. Dark grey profiles show depth of read coverage along reference sequence. Blue shading indicates spans where contigs aligned. The locations of spike and RNA-dependent RNA polymerase (RdRP) genes are indicated and shaded light grey.

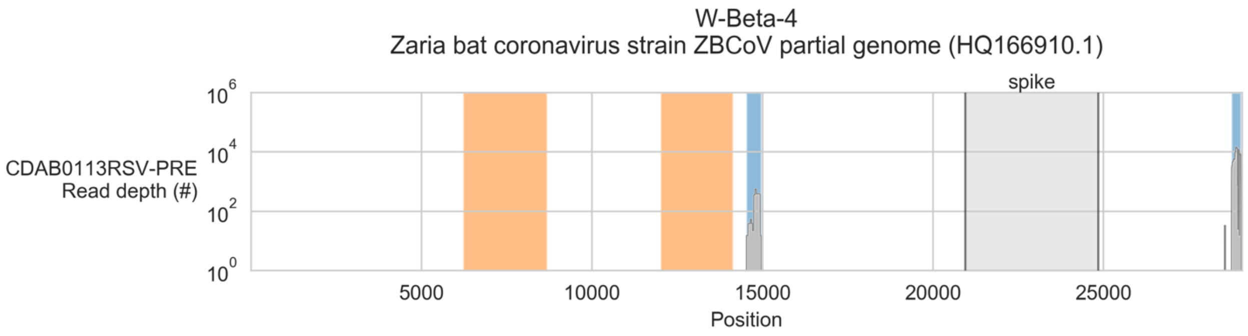


**Figure S4: Coverage of reference sequence by probe captured libraries for specimens from phylogenetic group W-Beta-4.** Coverage of reference sequence was determined by mapping reads and aligning contigs from probe captured libraries. Dark grey profiles show depth of read coverage along reference sequence. Blue shading indicates spans where contigs aligned. The locations of spike gene are indicated and shaded light grey. Ambiguous bases (Ns) are shaded orange.

### MATERIALS AND METHODS SUPPLEMENTAL

**Probe panel design:** All available sequences in the following taxa were downloaded from NCBI GenBank on October 4, 2020: unclassified *coronavirinae* (txid: 693995), unclassified *coronaviridae* (txid: 1986197), *alphacoronavirus* (txid: 693996), and *betacoronavirus* (txid: 694002). Bat CoV sequences were extracted by searching sequence headers for bat-related key words identified by the authors. These sequences were used as targets for probe design with the ProbeTools package (v0.0.5) (https://github.com/KevinKuchinski/ProbeTools) [Kuchinski 2022]. All possible probes were generated from the bat CoV sequences using the *makeprobes* module with a batch size of 100 probes. This generated a core panel of 18,365 probes. Since the next breakpoint in the manufacturer's pricing occurred at 20,000 probes, we designed additional probes targeting conserved motifs in CoVs from non-bat hosts. We used the *capture* and *getlowcov* modules to extract regions of the unclassified *coronavirinae*, unclassified *coronaviridae*, *alphacoronavirus*, and *betacoronavirus* sequences from all hosts not already covered by the core panel. These regions were then used as input targets for *makeprobes* with a batch size of 50 probes. The first 1,605 probes generated in this way became the supplemental panel.

While designing the supplemental panel, we removed SARS-CoV-2 sequences from the *betacoronavirus* space because they were over-represented and could have biased probe design towards this single taxon. To ensure coverage of SARS-CoV-2-related viruses by our panel, we used the *capture* and *getlowcov* modules to extract regions of the Wuhan-Hu-1 reference genome (MN908947.3) not already covered by the core panel. These regions were then used as input targets for *makeprobes* with a batch size of 1 probe, generating 29 probes that were added to the supplemental panel. The following were combined to create the final panel: the core panel of 18,365 probes generated from bat CoV sequences, the supplemental panel of 1,634 probes targeting conserved motifs in non-bat CoVs and SARS-CoV-2, and a single probe targeting our artificial control oligo sequence (see section on library construction and probe capture in main text's Materials and Methods). The final panel (Supplemental 2) was synthesized by Twist Bioscience (San Francisco, CA, USA).

**Library construction and pooling:** Sequencing libraries were constructed using the NEBNext Ultra II RNA Library Prep with Sample Purification Beads kit (E7775). 5 µl of undiluted RNA specimen was used as input for first strand synthesis. The fragmentation reaction incubation was shortened to 2 mins at 94 °C while the first strand synthesis incubations were modified to 10 mins at 25 °C, followed by 50 mins at 42 °C, followed by 10 mins at 70 °C. Second strand synthesis, bead clean-up, and end prep reactions were performed according to the kit's protocol. The adapter ligation incubation was extended to 60 mins at 20 °C, and the USER digest was also extended to 60 mins at 37 °C. Following another bead clean-up performed according to the kit protocol, libraries were barcoded with NEBNext Multiplex Oligos for Illumina (96 Unique Dual Index Primer Pairs) kit (#6440). Barcoding PCRs used the following cycling conditions: 1 cycle of 98 °C for 1 min; 12 cycles of 98 °C for 30 secs, then then 65 °C for 75 secs; 1 cycle of 65 °C for 10 mins. Barcoded libraries were purified with the final bead clean-up according to the kit's protocol.

**Probe capture:** Libraries were quantified with the Invitrogen Qubit dsDNA HS kit (Q32851), then 180 ng of each library was pooled together. The library pool was fully evaporated in a GeneVac miVac DNA concentrator (DNA-12060-C00) instrument. The dried library pool used to set up a hybridization reaction with 0.2 fmol/probe of our custom bat CoV probe panel (Twist Biosciences, San Francisco, CA, USA), Twist Universal Blockers (#100578), and the Twist Fast Hybridization Reagents kit (#101174) following the manufacturer's protocol. Hybridization reactions were incubated at 70 °C for 16 hours, then captured and washed with the Twist Binding and Purification Beads (#100983) and Twist Fast Hybridization Wash buffers (#101174) following the manufacturer's protocol until the final step, at which point the streptavidin bead slurry was resuspended in 22.5 µl of nuclease-free water instead of 50 µl. The entire 22.5 µl volume was used in the post-capture PCR, which was set up with NEBNext Ultra II Q5 2X Master Mix (#M0544), and Illumina amplification primers from the Twist Fast Hybridization Reagents kit (#101174). Post-capture PCRs were conducted with the following cycling conditions: 1 cycle of 98 °C for 60 secs; 25 cycles of 98 °C for 30 secs, then then 60 °C for 30 secs, then 65 °C for 75 secs; 1 cycle of 65 °C for 10 mins. Post-capture PCRs were purified using 0.8X SPRI beads from the Twist Binding and Purification Beads (#100983). Bead clean-up reactions were washed twice with 200 µl of 80% ethanol and eluted in 20 µl of nuclease-free water. Following the first capture, the captured pool was again completely evaporated, then a second capture was performed as before.

**Sequencing of captured libraries and removal of index hop artefacts:** Probe captured libraries were sequenced on an Illumina MiSeq instrument using V2 300 cycle reagent kits (#MS-102-2002). The double-captured library pool was sequenced across two MiSeq runs. The first run generated paired-end reads where each end was sequenced with 150 cycles. The second run generated paired-end reads where the first end was sequenced with 15 cycles and the second end was sequenced with 285 cycles. Index hops were filtered from both runs using HopDropper (v0.0.3) (https://github.com/KevinKuchinski/HopDropper) with UMIs of length 14, requiring a minimum base quality of PHRED 30, and discarding UMI pairs appearing only once. After removing index hops, reads from the second MiSeq run were treated as single-ended. This was done by discarding the short first end which was only necessary for index hop removal by HopDropper.

**De novo assembly of contigs from captured reads:** coronaSPAdes (v3.15.0) was used to assemble contigs *de novo* from probe captured MiSeq data [Meleshko 2021]. Reads from the first MiSeq run were provided to coronaSPAdes as paired-end data, while reads from the second MiSeq run were provided as single-end data. CoV contigs were identified using BLASTn (v2.5.0) against a local database composed of all *coronaviridae* sequences (txid: 11118) in GenBank available as of October 11, 2021 [Camacho 2009].

**Alignment of reads and contigs to bat CoV reference sequences:** Probe captured reads were mapped to selected reference genomes using bwa mem (0.7.17-r1188). Alignments were filtered with samtools view (v1.11) to retain properly paired reads (bitflag 3) and exclude unmapped reads, reads without mapped mates, not primary alignments, supplementary alignments, and reads failing platform/vendor quality checks (bitflag 2828) [Li 2009a, Li 2009b]. Samtools sort and index (v1.11) were then used to sort and index filtered alignments. Depth and extent of read coverage were determined with bedtools genomecov (v2.30.0) [Quinlan 2010]. Contig coverage was determined by aligning contigs to reference sequences with BLASTn (v2.5.0) and extracting subject start and subject end coordinates [Camacho 2009].

**Deep metagenomic sequencing of uncaptured libraries and generation of complete viral genomes:** New libraries were prepared from selected specimens following the same protocol as for libraries that were probe captured. These libraries were sequenced on an Illumina HiSeq X instrument by the Michael Smith Genome Sciences Centre (Vancouver, BC, Canada). Reads were assembled and scaffolded into draft genomes with coronaSPAdes (v3.15.3) [Meleshko 2021]. CoV-sized scaffolds were manually inspected to identify draft genomes. For one specimen (CDAB0492R), two contigs were manually joined to complete a complete draft genome.

HiSeq reads were mapped to draft genomes using bwa mem (v0.7.17-r1188). Alignments were filtered with samtools view (v1.11) to retain properly paired reads (bitflag 3) and exclude unmapped reads, reads without mapped mates, not primary alignments, supplementary alignments, and reads failing platform/vendor quality checks (bitflag 2828) [Li 2009a, Li 2009b]. Samtools sort and index (v1.11) were then used to sort and index filtered alignments. Variants were called with bcftools mpileup and call (v1.9) [Danecek 2021]. For bcftools mpileup, 30 was used as the minimum read mapping (-q) and base quality scores (-Q), and a minimum of 10 gapped reads was used for indel candidates (-M). For bcftools call, a ploidy of 1 was used (--ploidy). Low coverage positions in the draft genomes (less than 10 reads) were masked using bedtools genomecov (v2.30.0) [Quinlan 2010], then variants were applied to draft genomes with bcftools consensus (v1.9) to generate final complete genomes [Danecek 2021].

**Phylogenetic analysis of novel spike gene sequences:** Novel spike gene coding sequences were identified in three steps. First, we obtained the regions annotated as spike gene coding sequences from each study specimen's closest reference sequence in GenBank/RefSeq. Second, these spike coding sequences from the closest reference sequences were aligned to the final genomes of the novel bat CoVs using BLASTn (v2.5.0) [Camacho 2009]. Third, novel spike coding sequences were extracted using the subject start and end coordinates from the alignment. Novel spike CDSs were then translated using a custom Python script. Translated sequences were queried against all translated *coronaviridae* spike sequences in GenBank (available on October 11, 2021) using BLASTp (v2.5.0) [Camacho 2009]. For each genus, novel spike genes from study specimens were combined with the 25 closest-matching GenBank spike sequences (based on alignment bitscore) and all spike sequences available in RefSeq. Multiple sequence alignments were conducted with clustalw (v2.1) with default parameters, then phylogenetic trees were constructed from aligned sequences using PhyML (v3.3.20190909) with 100 bootstrap replicates [Thompson 1994, Guindon 2005].
